# Severely impaired bone material quality in *Chihuahua* zebrafish resembles classical dominant human osteogenesis imperfecta

**DOI:** 10.1101/251652

**Authors:** Imke A.K. Fiedler, Felix N. Schmidt, Christine Plumeyer, Petar Milovanovic, Roberta Gioia, Francesca Tonelli, Antonella Forlino, Björn Busse

**Author notes:** **Corresponding Author** Björn Busse, Ph.D., Department of Osteology and Biomechanics, University Medical Center Hamburg-Eppendorf, Lottestr. 55A, 22529 Hamburg, Germany, [**Disclosure** The authors have nothing to disclose.].

## Abstract

**Abstract:** Excessive skeletal deformations and brittle fractures in the vast majority of patients suffering from osteogenesis imperfecta (OI) are a result of substantially reduced bone quality. Since the mechanical competence of bone is dependent on the tissue characteristics at small length scales, it is of crucial importance to assess how osteogenesis imperfecta manifests at the micro- and nanoscale of bone. In this context, the *Chihuahua* (*Chi/+*) zebrafish, carrying a heterozygous glycine substitution in the α1 chain of collagen type I, has recently been proposed as suitable animal model of dominant OI. Similar to human severe OI type III, *Chi/+* show skeletal deformities, altered mineralization patterns and a smaller body size. Using a multimodal approach targeting bone quality parameters, this study aims at quantifying the changes in bone morphology, structure and tissue composition of *Chi/+* at multiple length scales. Morphological changes were assessed with high-resolution micro-CT imaging and showed that the vertebrae in *Chi/+* had a significantly smaller size, thinner cortical shell and distorted shape. Tissue composition in vertebrae was investigated with quantitative backscattered electron microscopy and Fourier-transform infrared spectroscopy, showing higher mean calcium content, greater matrix porosity, as well as lower mineral crystallinity and collagen maturity in comparison to controls. This study provides comprehensive quantitative data on bone quality indices in *Chi/+* and thus further validates this mutant as an important model reflecting osseous characteristics associated with human classical dominant osteogenesis imperfecta, both at the whole bone level and the tissue level.

## Introduction

Osteogenesis imperfecta (OI) is a genetic disorder of bone and connective tissues which affects approximately 1 in 10,000 births and has multiple implications on skeletal growth and bone mineralization ^(1)^. The mechanical integrity of the diseased bone is often drastically impaired, leading to an increased bone fragility ^(2,3)^. As a consequence, patients suffer from excessive skeletal deformations and multiple brittle fractures ^(3–5)^. The compromised fracture resistance in the vast majority of OI patients is due to a dominant mutation of the genes responsible for synthesis of collagen I, *i.e*. the organic framework of bone ^(6–8)^. This results in a diminished structure of the organic matrix accompanied by delayed mineralization and thus retardation in skeletal growth and maturation.

The severity of the disease can vary largely and depends on the type of OI^(9)^. Among the dominant mutations affecting the collagen I genes, the mildest and most common form of OI is type I with spontaneous fractures occurring after a child started walking, minor deformities, an almost normal stature, and mild bone fragility. The most severe and lethal form is type II with fractures occurring *in utero* and death before birth. Intermediate forms are type III with severe phenotypes including fractures, scoliosis, major deformities and a very small stature, and type IV with moderate phenotypes including fractures and a small stature ^(10)^.

Since these types of OI often evidently manifest at the skeletal and whole bone level in terms of low bone mass and the presence of fractures or deformities, initial diagnostic approaches commonly employ X-ray absorptiometry and radiography ^(1,7,11)^. Based on bone biopsies from patients, OI has further been characterized at the tissue level, *i.e.* at the micro-and nanoscale of bone, where the ultrastructural alterations have been observed to correlate well with the clinical severity of OI ^(12)^. For tissue of OI type I to IV there have been reports on impaired mechanical behavior at sub-micron scale ^(13,14)^, the presence of irregularly hyper-mineralized regions ^(14)^ and changes in the structure of the nanoscopic bone components, *i.e.* collagen fibrils and hydroxyapatite crystals ^(14)^. Since the material properties of bone at small length scales, *i.e.* bone tissue quality, determines the mechanical competence of bone at the whole bone and skeletal level, it is of crucial importance to assess in which way the bone tissue quality is compromised by OI ^(13)^. Yet, current treatment options are mainly aiming at increasing bone strength through increasing overall bone mass ^(15–17)^. Conventional measures including musculoskeletal exercise are therefore often combined with pharmacological approaches widely based on antiresorptive bisphosphonate therapy ^(8,15)^. However, given the particularly wide age range of patients in addition to the large clinical variability and the genetic diversity of OI, the success of such therapies often remains inconclusive ^(18,19)^.

Thus, for better understanding the impact of the disease on all skeletal length scales, and to accelerate the search for enhanced treatment options of the brittle bone disease, a large number of animal models have been established and characterized ^(20,21)^. Besides well studied mammalian models including mice ^(22–25)^ and dogs ^(26)^, novel small-sized animal models are emerging. The zebrafish *(Danio rerio)* has become a valued animal model in the field of biomedical research due to its short generation time, genetic similarity to humans, small size, and low husbandry costs ^(27)^. In the context of OI and similar genetic bone disorders, few zebrafish mutations have so far been generated and characterized with regard to the skeletal phenotype ^(21,28,29)^. Fisher *et al.* have first described the dominant G574D mutation in the *Chihuahua* (*Chi/+*) zebrafish, which affects the *col1a1a* gene encoding for α1 chain of collagen I and therefore shares genetic similarities to severe dominant human OI. Carrying a glycine substitution in the α1(I), these zebrafish suffer from defects and retardation in bone growth and maturation ^(30)^. Recently, this mutant has been characterized more deeply by Gioia *et al* ^(31)^, where static and dynamic bone formation markers the have confirmed severe skeletal deformities and delayed mineralization, confirming the suitability of *Chi/+* mutants as promising model for classical dominant OI.

To expand the validity of *Chi/+* as model for human OI, further data on the material properties influencing the fracture behavior of bone is needed. Using a multimodal array of techniques focusing on the macro- to nanoscale of zebrafish bone, the study aims to provide quantitative data on the alterations in bone morphology, structure and composition in *Chi/+* zebrafish from the skeletal to the tissue level.

## Materials & Methods

*Chihuahua* (*Chi/+*, *col1a1*a^dc124/+^) and wild-type AB (WT) zebrafish were bred in-house at the animal facility “Centro di servizio per la gestione unificata delle attività di stabulazione e di radiobiologia” of the University of Pavia. The mutant *Chi/+* carries a G2207A mutation in *col1a1a*, causing a heterozygous G574D substitution in the α1 chain of type I collagen ^(30)^. Husbandry, breeding and housing conditions of the zebrafish are described in detail in Gioja *et al.* ^(31)^. All animals have been sacrificed at 10 months and kept frozen at −80° C (n=12/group).

### Micro-Computed Tomography (Micro-CT)

Skeletal morphology and bone microstructures were assessed using micro-CT at a resolution of 5 μm (Skyscan 1272, Bruker, Kontich, Belgium). Six fish per group were fixed in 3.5 % formalin for 24 h and placed in a moist chamber during scanning. Whole body scans were acquired at 55 kV and 166 μA without an X-ray filter. Ring artifact and beam hardening correction was kept constant for all samples during reconstruction with NRecon (Bruker, Kontich, Belgium). After applying a fixed threshold for all samples, 3D evaluation was conducted using CTAn (Bruker, Kontich, Belgium). Quantification of changes in bone morphology of *Chi/+* and WT was performed on the first and second precaudal vertebrae of each fish. Hereby, neural and hemal arches were excluded from the analysis. Determined parameters were bone volume (BV, mm^3^), defined as the vertebral body without the inner space; vertebral body height (VBH, μm); vertebral thickness (V.Th, μm); and eccentricity (Ecc, 0-1).

### Fourier-Transform Infrared (FTIR) Spectroscopy

The molecular composition, maturity of the mineral- and collagen phase as well as matrix porosity was evaluated by FTIR spectroscopy. Six zebrafish per group were fixed in 3.5 % formalin for 24 hours, subsequently underwent step-wise dehydration in increasing concentrations of ethanol, and were embedded in poly methyl methacrylate (PMMA). Sample blocks were polished coplanar for tissue composition analysis, whereby the precaudal vertebral end plates cut in lateral plane were selected as region of interest. *Chi/+* and WT vertebrae were scanned using FTIR spectroscopy (Spotlight 400 attached to Frontier 400, PerkinElmer, Waltham, MA, USA) in Attenuated Total Reflectance (ATR) mode. Maps with dimensions of 65 μm × 130 μm were acquired at a spatial resolution (*i.e.* pixel size) of 1.56 μm with eight scans per pixel and a wavenumber range from 4000 – 570 cm^- 1^. Post-processing and spectral analysis was performed using a custom-written Matlab script (R2016b, MathWorks, Natick, MA, USA) and included the subtraction of the PMMA signal, smoothing of the absorbance signal and linear baseline removal. Peaks of interest were identified based on the second derivative of the original spectra. The phosphate peak was located between 1154 and 900 cm^- 1^, the carbonate peak was centered at 870 cm^-1^, and the amide I peak envelope was located between 1710 and 1600 cm^−1^. Bone tissue parameters were determined by dividing respective peak areas, including the mineral-to-matrix ratio (*i.e.* area of phosphate/amide I), carbonate-to-phosphate ratio (*i.e.* area of carbonate/phosphate), crystallinity (*i.e.* area of phosphate sub peaks 1028/1018 cm^-1^), and collagen maturity (*i.e.* amide I sub peaks 1660/1690 cm^-1^. Matrix porosity was determined by relating the PMMA peak located at 1725 cm^-1^to the amide I peak. Using Raman spectroscopy with a spatial resolution of < 1 μm, this approach has recently been used to determine the parameter nano-porosity^(32)^. Due to the lower resolution of 1.56 μm used in this present study, the parameter also includes porous structures above the nanometer scale. Hence, the parameter is hereinafter referred to as matrix porosity.

### Quantitative Backscattered Electron Imaging (qBEI)

For analyzing the bone mineral density distribution (BMDD) in precaudal vertebrae of *Chi/+* and WT zebrafish, electron imaging was performed on identical samples and in identical regions of interest as previously analyzed by FTIR spectroscopy. Prior to imaging, samples were carbon-coated. Six specimens per group were investigated using a scanning electron microscope (LEO 435 VP; LEO Electron Microscopy Ltd., Cambridge, UK) operated at 20 kV and 680 pA using a constant working distance of 20 mm (BSE Detector, Type 202; K.E. Developments Ltd., Cambridge, UK). BMDD was determined on grey value images according to previously described protocols ^(33–35)^. Calibration of the system was performed using a carbon-aluminum standard, facilitating the quantification of bone mineral as calcium weight percentage. Determined parameters were the mean calcium content (CaMean, wt%), the heterogeneity of the bone mineral distribution (CaWidth, wt%), the amount of low mineralized bone (CaLow, %), and the amount of highly mineralized bone (CaHigh, %).

### Statistical Analysis

Statistical analysis was performed using SPSS (Version 24, IBM, Armonk, NY, USA). Normal distribution and homogeneity of variance was tested using Shapiro-Wilk tests and Levene’s test, respectively. Group comparisons were carried out using independent t-tests at a significance level of α=0.05. Results are presented as mean±one-fold standard deviation.

## Results

Bone quality analyses performed on *Chi/+* and their age-matched WT controls gained distinct bone phenotypes of the mutation at several length scales. At the skeletal level, predominant manifestations included a shorter body length and abnormal mineralization patterns. At the whole bone level, precaudal vertebrae of *Chi/+* showed substantially altered morphology. At the tissue level, vertebrae of *Chi/+* presented a high degree of mineralization, low mineral crystallinity and low matrix maturity in addition to elevated matrix porosity.

### Skeletal morphology and whole bone structure

Micro-CT 3D reconstructions of *Chi/+* and WT zebrafish revealed substantial differences between the groups regarding their macroscopic skeletal morphology and mineralization (Fig. 1 A, B). *Chi/+* zebrafish had significantly reduced body length compared to WT with 17.1±2.8 mm vs. 27.5±1.8 mm (p=0.001). Along with accumulated fractures and deformations in the anal, dorsal and caudal fin bones, in the ribs, and in the neural and hemal arches of the vertebrae, the skeletal apparatus of *Chi/+* zebrafish presented heterogeneous mineralization as indicated by differences in X-ray absorption. At the whole bone level, particularly along the spine of *Chi/+* zebrafish, regions of high mineralization were observed directly adjacent to regions of lower mineralization, *i.e.* highly mineralized and fused vertebrae in precaudal and caudal regions were located next to low mineralized spinal regions in-between.

**Figure 1.**
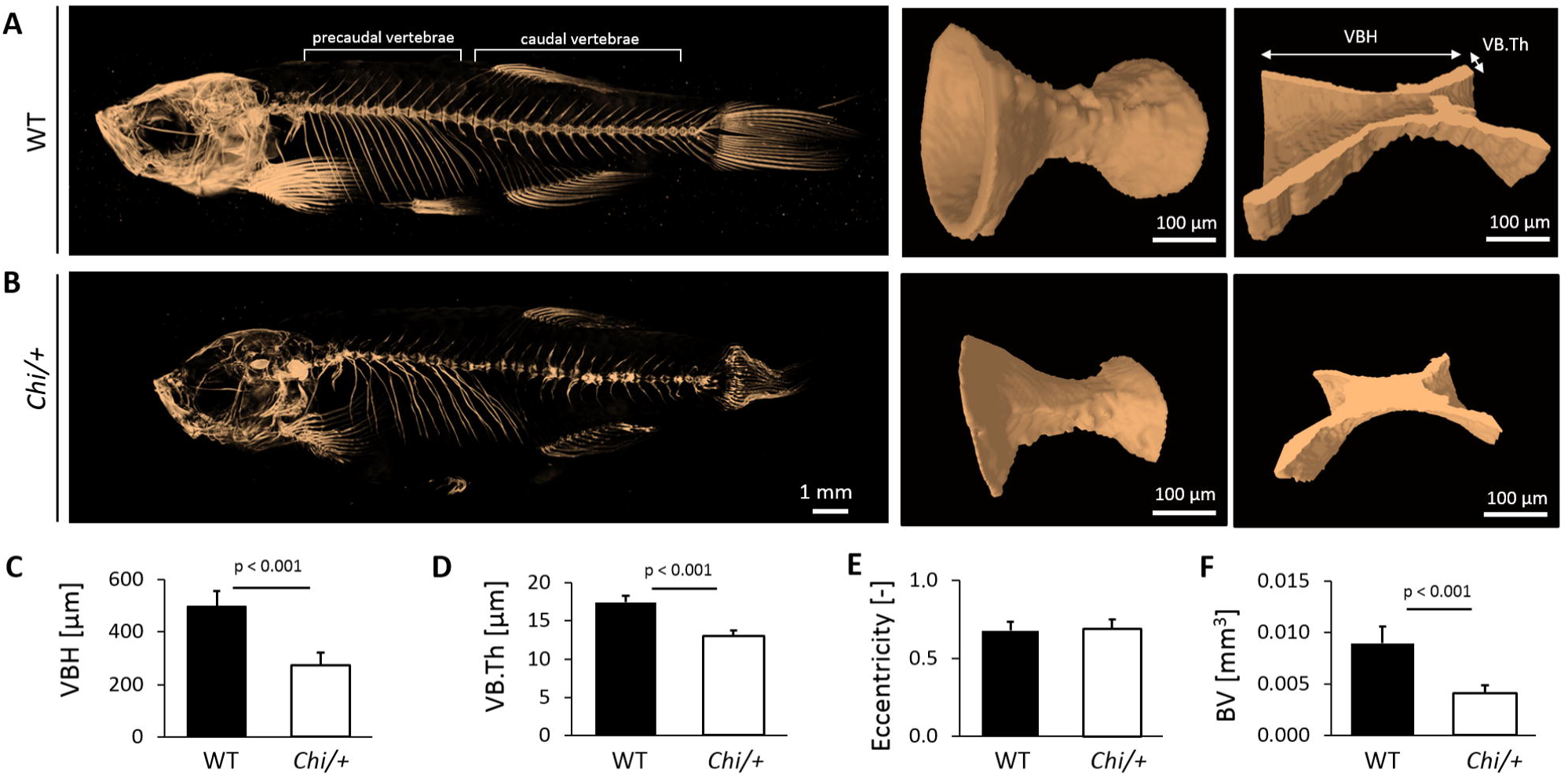
Skeletal morphology and structure of *Chihuahua* and WT zebrafish vertebrae based on micro-CT.**A)** Representative whole body scan of an adult wild type zebrafish (left) and an isolated precaudal vertebral body with typical double-cone geometry (center). Histomorphometric parameters are indicated in the longitudinal section of the vertebral body (right). **B)** Representative whole body scan of a *Chi/+* mutant (left) indicates heterogeneous mineralization in the skull and along the spine, deformities of ribs and fins possibly related to fractures, and fused or collapsed vertebrae. Vertebral bodies of *Chi/+* mutants showed abnormal geometry (center) and accumulation of mineralized tissue in its core (right). Histomorphometric analysis of precaudal vertebrae yielded **C)** a significantly smaller vertebral body height in *Chi/+*, **D)** significantly reduced vertebral thickness, **E)** similar eccentricity, and **F)** reduced bone volume. Based on skeletal features and whole bone characteristics, the *Chihuahua* zebrafish clearly resembles human severe osteogenesis imperfecta type III.

The quantification of bone morphological differences in the precaudal vertebrae gained significantly smaller vertebral bodies in *Chi/+* compared to WT zebrafish with a significantly lower vertebral body height compared to WT with 288±44 μm vs. 488±52 μm, p<0.001 (Fig. 1 C), as well as significantly lower vertebral thickness with 13.5±1.0 μm vs. 17.2±0.8 μm, p<0.001 (Fig. 1 D). Notably, precaudal vertebral bodies of *Chi/+* presented a mineralized core (Fig. 1 B, right), which was excluded from the morphometric evaluation. While the eccentricity (*i.e.* roundness) of vertebral bodies was similar between both groups (Fig. 1 E), *Chi/+* zebrafish presented a significantly reduced BV of 0.004±0.001 mm^3^ vs 0.008±0.001 mm^3^ in WT controls, p<0.001 (Fig. 1 F).

### Tissue composition, maturity and porosity

Bone tissue composition analysis of *Chi/+* and WT zebrafish vertebrae by FTIR spectroscopy gained substantial differences in the majority of mineral- and collagen-related parameters (see Fig. 2 A-E). The carbonate-to-phosphate ratio was significantly higher in *Chi/+* compared to WT with 0.092±0.002 vs. 0.089±0.002, p<0.01 (Fig. 2 A), while the parameter crystallinity, a measure for crystal maturity and size, was significantly lower in *Chi/+* zebrafish compared to WT controls with 11.5±1.6 vs. 20.1±6.3, p=0.005 (Fig. 2 B). The mineral-to-matrix ratio was insignificantly elevated in *Chi/+* (Fig. 2 C). In addition, the collagen maturity ratio was significantly lower in *Chi/+* compared to WT with 10.1±5.8 vs. 24.2±5.7, p=0.003 (Fig. 2 D). With regard to the matrix porosity of vertebral tissue, *Chi/+* presented significantly higher values than WT with 0.184±0.085 vs. 0.090±0.022, p = 0.025 (Fig. 2 E).

**Figure 2.**
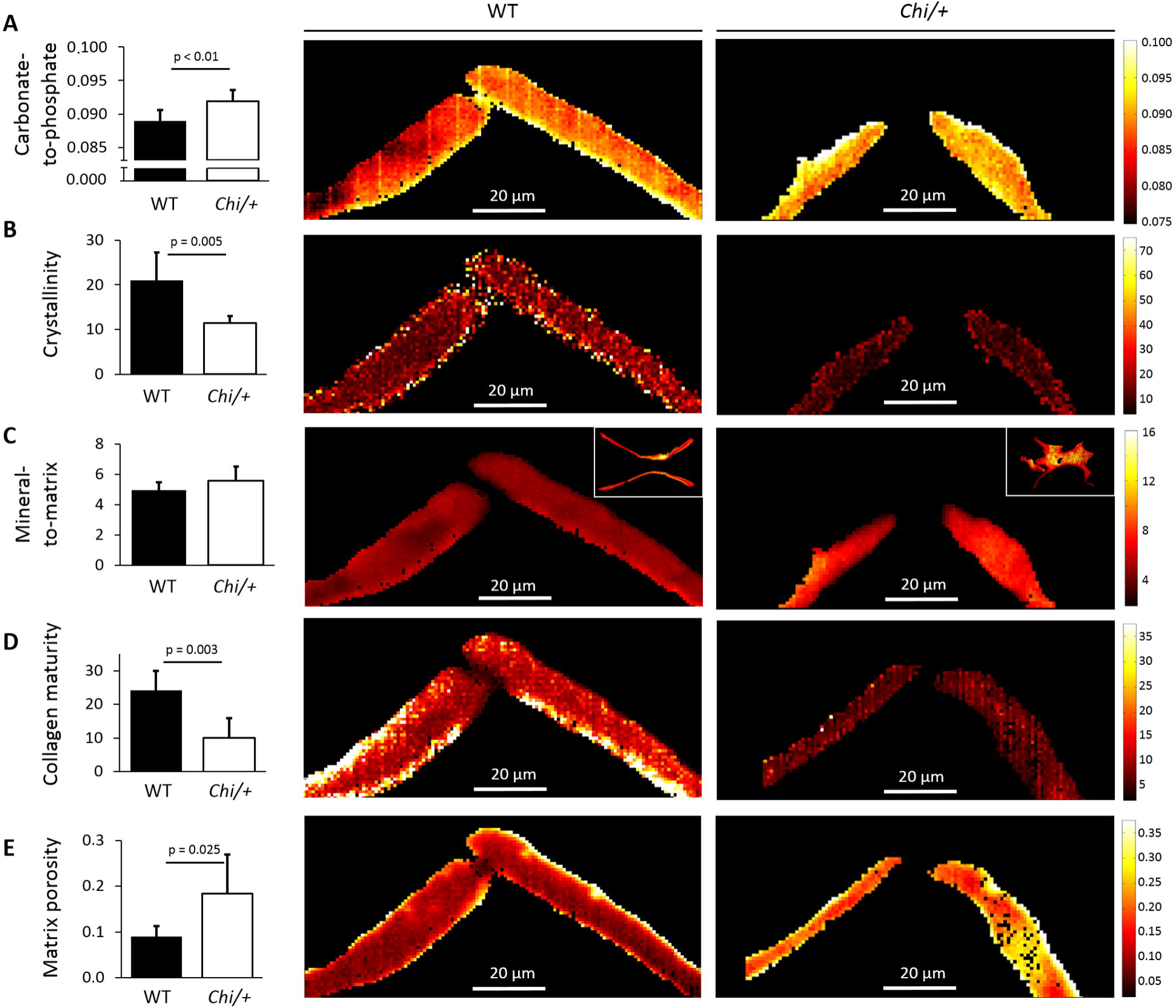
Molecular composition and structure assessed in precaudal vertebrae of WT and *Chi/+* using FTIR spectroscopy **A)** The carbonate-to-phosphate ratio, a measure for carbonate substitution in the crystal lattice, was significantly higher in *Chi/+*. **B)** Crystallinity, a measure for mineral crystal size, lattice purity and crystal age, was significantly lower in *Chi/+* mutants. **C)** The average mineral-to-matrix ratio was insignificantly higher in *Chi/+* than in WT. The inserts show representative spectral maps from whole vertebrae. Note the typical double-cone shape in WT in contrast to an altered structure in *Chi/+* including a mineralized core in *Chi/+*, which has a high mineral-to-matrix ratio (yellow pixels) in comparison to the cortical shell in both WT and *Chi/+* (red pixels). **D)** Collagen maturity was substantially reduced in *Chi/+* mutants. **E)** Matrix porosity was significantly elevated in *Chi/+* mutants compared to WT. Tissue composition analysis revealed clear implications of the mutation on the tissue quality in zebrafish which are characteristic for classical dominant osteogenesis imperfecta.

### Bone mineral density distribution

BMDD acquired with quantitative backscattered electron microscopy showed clear differences in the mineralization pattern between *Chi/+* and WT controls (Fig. 3 A-E). As depicted in Fig. 3 B, the distribution of calcium was shifted towards higher concentrations in *Chi/+*. The mean calcium weight percentage of both groups was slightly below 30 wt%, whereby CaMean of *Chi/+* was significantly higher compared to WT with 27.8±0.8 wt% vs. 26.7±0.9 wt%, p=0.039 (Fig. 3 C). At tissue level, the heterogeneity of mineralization measured by CaWidth was similar between the groups as depicted in Fig. 3 D, and so was the percentage of low calcium concentrations CaLow. High calcium concentrations CaHigh in *Chi/+* were twofold compared to WT with 25.1±12.9% vs. 13.5±6.4 %, p=0.075 (Fig. 3 E).

**Figure 3.**
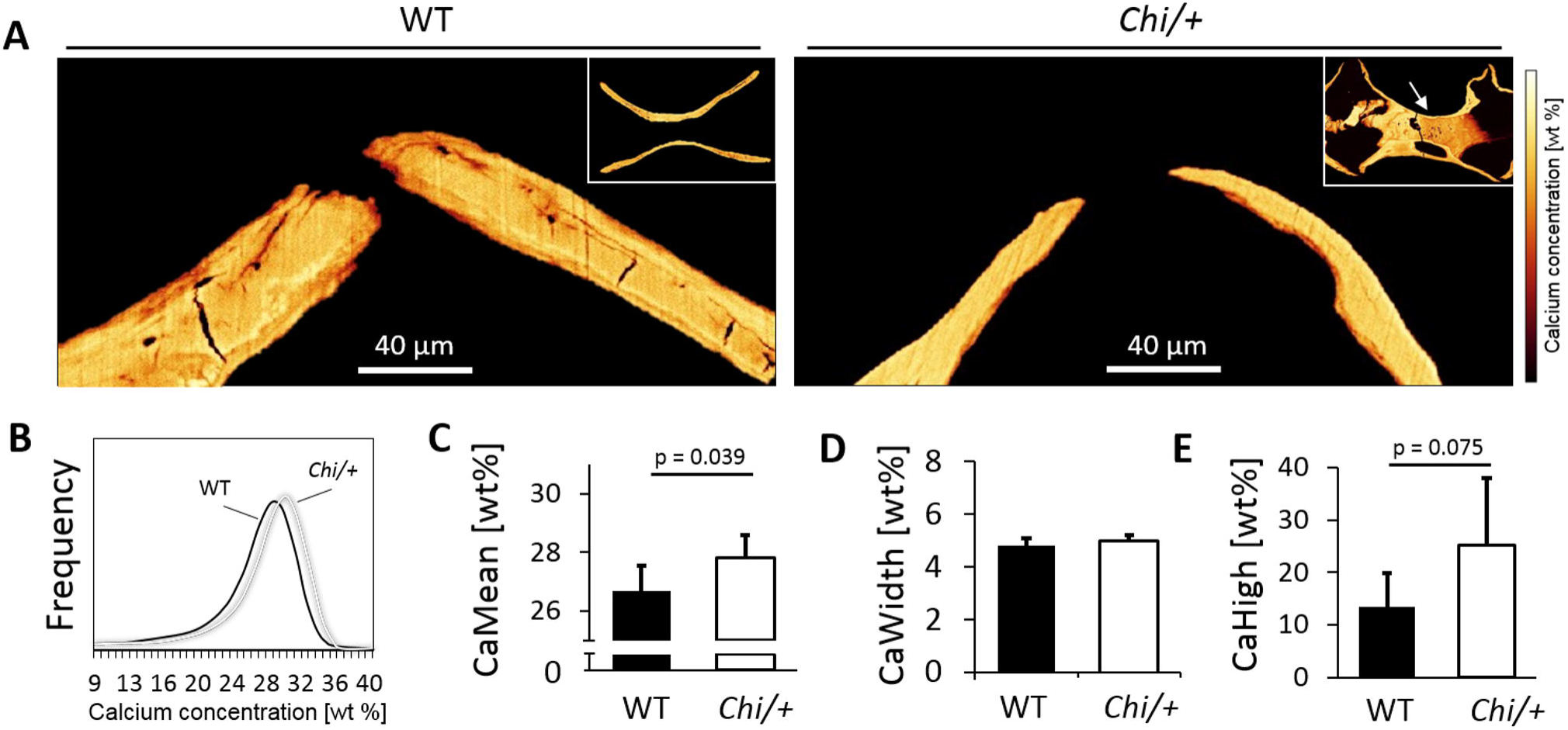
Bone mineral density distribution of WT and *Chi/+* zebrafish. **A)** The vertebral bodies yielded substantial differences in calcium concentration and distribution with brighter gray values in *Chi/+* than in WT. Inserts show representative qBEI images from whole precaudal vertebrae depicting a heterogeneously and lower mineralized core compared to the cortical shell (darker pixels) in *Chi/+* (arrow). Coupled with a high mineral-to-matrix ratio detected in the core tissue compared to the shell (Fig. 2 A), this tissue appears of lower degree of mineralization and lower collagen content compared to the cortical shell. **B)** The calcium concentration histogram was shifted towards higher concentrations in the *Chi/+* zebrafish. **C)** At tissue level, the mean calcium concentration was significantly elevated in *Chi/+* compared to WT. **D)** The heterogeneity of the calcium distribution was similar between the groups. **E)** Averaged high calcium concentrations were twofold in *Chi/+* compared to WT. The assessment of bone mineralization at tissue level shows substantial similarities to observations made in classical dominant OI.

## Discussion

Applying an array of bone quality analyses including micro-CT, qBEI and FTIR spectroscopy this study revealed and quantified changes in the morphology, structure and molecular composition of bone from the osteogenesis imperfecta zebrafish model *Chihuahua* carrying a heterozygous glycine substitution in the α1 chain of type I collagen. Alterations in bone phenotype induced by the mutation manifested at all investigated skeletal length scales, *i.e.* from skeletal to whole bone to tissue level.

3D histomorphometry performed on whole skeleton of *Chi/+* and WT zebrafish validated the presence of skeletal deformities and the occurrence of fractures described in recent radiographic and histological investigations performed on this *Chi/+* zebrafish model ^(31)^. Similar to cases of human OI, these abnormalities are present in both the axial and the appendicular skeleton ^(5)^. Focusing on the vertebral bodies at whole bone scale, results of this study showed substantially decreased bone volume and cortical shell of the vertebrae, which goes well in line with clinical observations on human osteogenesis imperfecta cases type I-IV ^(3,36,37)^. Interestingly, the eccentricity of the vertebral bodies, *i.e.* their roundness, was not affected by the mutation, indicating that the lateral locomotion patterns of *Chi/+* zebrafish did not lead to permanent malformations at whole bone level along the spine. Yet, *Chi/+* zebrafish presented distinct vertebral features visible in the first precaudal and mid-caudal vertebrae, which enclosed an atypical mineralized core. The mineral-to-matrix ratio distribution in whole precaudal vertebrae (Fig. 2 A, inserts) indicates a high ratio of phosphate to amide I inside this mineralized core (bright pixels) when compared to cortical vertebral regions (darker pixels). This indicates either a high degree of mineralization or a low degree of collagen content within this region. qBEI images of the identical region in the precaudal vertebrae (Fig. 3 A, inserts) showed highly heterogeneous mineral distribution in the core and low calcium content compared to cortical regions (low gray values). Combining these results, mineralized tissue in the core of *Chi/+* vertebrae seems to be a heterogeneous, low mineralized tissue with low presence of collagen.

In cases of human OI, the accumulation of multiple vertebral fractures has been identified to cause progressive spinal deformity ^(38)^, thus it is likely that the observed deformations along the spinal column of the *Chi/+* zebrafish could also be linked to a history of fractures. In a clinical setting, substantial macroscopic anomalies including fractures and deformities often allow the identification of OI based on radiographic indications ^(1)^. However, this approach is not sufficiently conclusive to also serve as basis for reliable fracture risk assessment, since the mechanical competence of healthy and pathological bone is determined by the material properties at tissue level ^(13)^. In this context, mouse models of OI have considerably aided in correlating the mechanical properties of OI-affected bone with ultra-structural defects at tissue level, and thus in better understanding the origins of OI-related bone fragility ^(24,39–42)^.

In this study on zebrafish carrying the α1-G574D mutation, the tissue composition analysis revealed that both the organic collagen matrix and the inorganic mineral crystal phase were significantly affected in *Chi/+* vertebral bone. While at skeletal level OI-affected bones present with reduced bone mineral density and bone mass ^(6)^, the mineralization pattern of OI can be highly heterogeneous throughout the skeleton. To that effect, an increased degree of mineralization at tissue level has been reported for murine OI models ^(24,39,42)^ and human OI ^(43–45)^. Analogous, the mineral density distribution in the bone tissue of the *Chi/+* zebrafish was shifted towards higher calcium concentrations compared to WT zebrafish. Hence, *Chi/+* zebrafish present with elevated mean calcium concentrations, while in contrast, the mineral crystallinity of the hydroxyapatite particles was significantly decreased. The parameter crystallinity as assessed with vibrational spectroscopy has been correlated with the maturity, size and purity of the hydroxyapatite crystals in bone, and thus indicates smaller, less stoichiometric crystals in *Chi/+* zebrafish. Moreover, *Chi/+* zebrafish showed a high carbonate-to-phosphate ratio, supporting high carbonate substitution in the crystal lattice composition and thus lower crystal maturity.

These findings in zebrafish are well in line with previous observations made in murine OI models ^(46)^, and in human tissue of OI type I-IV, suggesting that affected tissue comprises an increased mineral-to-matrix ratio and tissue mineral density (TMD) with smaller but more abundant crystals ^(13,43)^. In addition to a smaller size of crystals, the orientation of crystals in an OI-mouse model have been shown to more random than in healthy cases ^(47)^. While this observation is most probably linked to a more random orientation of collagen fibrils, it suggests an additional parameter of interest to investigate in the bone tissue of *Chi/+* mutants.

In addition to altered mineral properties, the molecular structure of the collagen matrix differed significantly in *Chi/+* zebrafish as evidenced by vibrational spectroscopy. While it should be noted, that the collagen-related sub peak positions and ratios employed in the spectral analysis have not yet been correlated to the collagen I molecule of zebrafish bone using biochemical analyses (*e.g.* with high performance liquid chromatography), we found that the established ratio for collagen maturity (1660/1690 cm^-1^) ^(48)^ yielded a significant difference towards lower collagen maturity in *Chi/+*. In combination with reduced crystallinity, these parameters may relate to an increased remodeling rate, which has been described in cases of human OI ^(37,43,49)^. The amount of low calcium values determined with quantitative backscattered electron microscopy has been correlated to the amount of bone which undergoes primary mineralization ^(43)^. As this value was not altered in *Chi/+* zebrafish, it stands to reason that the turnover kinetics in this zebrafish model are similar to human OI cases, featuring increased resorption rates at normal bone formation rates ^(43)^.

Besides compositional changes of the collagen and mineral phase, the tissue of *Chi/+* mutants further showed a higher matrix porosity, indicating substantial alterations in the microscopic structure. Hereby, the parameter matrix porosity was determined as the ratio of PMMA to amide I, as previously derived with Raman spectroscopy ^(32)^. During the tissue preparation process, the latter infiltrates into nano- and microscopic spaces that are void of mineral and collagen, *e.g.* within the crystal lattice or within the inter- and intra-fibrillar space. Based on Raman spectroscopic analyses using this parameter with sub-microscopic resolution on biopsies from children suffering from OI compared to healthy controls, it was recently suggested, that smaller but more abundant and densely packed crystals in OI cases are leading to a decrease in nanoporosity ^(32)^. However, this decrease has been found in immature tissue, *i.e.* in mineralization fronts of recently remodeled tissue (1-3 days), but not in more mature tissue. Based on observations made in classical dominant OI cases ^(50)^, bone tissue of *Chi/+* zebrafish likely features more randomly organized collagen fibrils with more inter-fibrillar cavities, offering an explanation for the herein detected elevated matrix porosity above the micron scale. Such micro-structural changes coupled with an altered chemical composition of the collagen and mineral components compromise the mechanical properties of the tissue, consequently leading to an increased bone fragility characteristic for classical dominant OI.

This study provides quantitative data on the bone morphology, structure and tissue composition of zebrafish carrying a dominant collagen mutation. Multimodal and multiscale bone characterization revealed a drastic impairment of bone quality in the skeleton of *Chi/+* zebrafish. Specific phenotypes at the skeletal level include a shorter skeletal size, accumulation of bone deformities and heterogeneous mineralization patterns, and at tissue level an increased mineral density, reduced mineral and matrix maturity, and increased matrix porosity. Detected at all investigated length scales, these characteristics are clearly resembling to human OI. Analogous to human OI cases, where the mechanical properties of bone are impaired, the observed deformities in *Chi/+* mutants are a consequence of altered bone quality at tissue level. The *Chihuahua* zebrafish can therefore be expected to provide an exceeding animal model for human OI. Further investigations of this model will be of great value for improving current treatment strategies of OI, particularly in the form of novel drug screening tests and drug administration as well as in the form of conventional therapies including musculoskeletal exercise.

## Acknowledgements

The authors thank Dr. Shannon Fisher (Department of Pharmacology & Experimental Therapeutics, Boston University School of Medicine, USA) for providing the mutation of *Chihuahua* zebrafish and Dr. Katharina Jähn (Department for Osteology and Biomechanics, University Medical Center Hamburg-Eppendorf) for her assistance. The research leading to these results received funding from PIER (Partnership for Innovation, Education and Research) under grant no. PIF-2014-28 and DFG (Deutsche Forschungsgemeinschaft/ German Research Foundation) under grant no. BU 2562/2-1/3-1 to BB; from Fondazione Cariplo (grant No. 2013-0612) and Telethon (grant No. GGP13098) to AF; and from the Joachim Herz Stiftung in cooperation with PIER to FNS.

### Authors’ contributions

IAKF, PM, AF and BB designed the study; RG, FT and AF generated the fish for this study; IAKF, PM, FNS and CP prepared the specimens for experimental analyses; IAKF, FNS, CP, and PM acquired and analyzed the data; IAKF performed the statistical analysis; IAKF, FNS, PM, RG, FT, AF and BB interpreted the data. IAKF and BB wrote the manuscript. All authors approved the final version of the manuscript.

